# Genome characterization of monkeypox cases detected in India: Identification of three sub clusters among A.2 lineage

**DOI:** 10.1101/2022.09.16.507742

**Authors:** Anita M. Shete, Pragya D. Yadav, Abhinendra Kumar, Savita Patil, Deepak Y. Patil, Yash Joshi, Triparna Majumdar, Vineet Relhan, Rima R. Sahay, Meenakshy Vasu, Pranita Gawande, Ajay Verma, Arbind Kumar, Shivram Dhakad, Anukumar Bala Krishnan, Shubin Chenayil, Suresh Kumar, Priya Abraham

**Affiliations:** Indian Council of Medical Research-National Institute of Virology, Pune, Maharashtra, India, Pin-411021; Maulana Azad Medical College and Lok Nayak Hospital, New Delhi, India, Pin-110002; Public Health Department of Kerala, Directorate of Health Services, Thiruvananthapuram, India, Pin - 695 035; All India Institute of Medical Sciences, New Delhi, India, Pin-110029; Indian Council of Medical Research-National Institute of Virology, Alappuzha, Kerala, India, Pin-688005; State Surveillance Unit (IDSP), Directorate of Health Services (IDSP), Malappuram, Kerala, India, Pin-676505

**Keywords:** Monkeypox cases, Genomic characterization, Phylogeny, India, APOBEC3, SNP

## Abstract

Since May 2022, Monkeypox, a zoonotic *Orthopox* DNA virus was reported in more than 102 countries indicating expansion of its geographic range. We analyzed the complete genomes sequences of Monkeypox cases from Kerala (n=5 travelled from UAE) and Delhi (n=5 no travel history), India confirmed during July to August 2022. All the retrieved MPXV sequences from India covering 90 to 99% genome belong to A.2 lineage of clade IIb. The A.2 MPXV lineage divided in three sub clusters; first cluster Kerala n=5, Delhi n=2 aligned with the USA-2022 ON674051.1; while second of Delhi n=3 aligned with USA-2022 ON675438.1 and third consists of the UK, USA and Thailand. Recent update in MPXV lineage designated all the five sequences from Kerala as A.2.1. In addition to known 16 single nucleotide polymorphisms (SNPs) along with 13 APOBEC3 cytosine deaminase activity determined specific lineage defining mutations in A.2 lineage, 25 additional APOBEC3 mutations were found in 10 reported sequences. The study emphasizes need of enhancing genomic surveillance to understand the mutation and its linkage.

## Introduction

Monkeypox virus (MPXV) belongs to the genus *Orthopoxvirus* which causes infection in humans. The genus also includes the dreadful variola virus (VAV) which causes smallpox, cowpox virus, camelpox virus, and vaccinia virus.^1^ Although the clinical features of the monkeypox closely resemble the smallpox, MPXV is not ancestral to the VAV.^2^ The MPXV is considered to be one of the largest genomes of 197 kb and it is the most complex virus.^3^ The first human case was reported in 1970 from the Democratic Republic of Congo. Since then, it has spread to central and West African countries and became endemic to these countries. Outside Africa, United States reported the first Monkeypox outbreak in the year in 2003. The United Kingdom (UK) reported its first Monkeypox cases in September 2018.^4^ The sporadic cases of Monkeypox were reported around the world in travellers arriving from Nigeria, Britain, Israel and Singapore.5,6 Recently, massive outbreaks of Monkeypox have been observed in 102 locations affecting 60,799 confirmed cases and 20 deaths from both endemic and non endemic countries.7 As per the recent genome classification of the World Health Organization, the MPXV genome is now designated as Clade I (Congo Basin clade), Clade II (West African clade) with sub-clades Clade IIa and Clade IIb. The MPXV strain belonging to Clade IIb has been the cause of recent 2022 Monkeypox outbreak.^8^

MPXV is a double stranded DNA genome that includes a core region, a left arm, and a right arm which encodes about ∼ 190 genes. Core region encodes a conserved region for viral replication and assembly genes are encoded by the core region. Inverted terminal repeats (ITR) are located in the left and right variable regions which lays important role in the pathogenicity of the Monkeypox virus.^9^ Mainly the highly conserved genes responsible for housekeeping functions are present in the central region of the genome while the genes encoding for the virus–host interactions and are located in the terminal region.^10^ High recombination frequencies were noted in *Poxviridae* family with inter and intra species recombination which might cause gain or loss of genetic material.^11^ The difference of 900bp has been documented in West Africa (WA) and the Congo Basin (CB) lineage.^12^ Till now a low mutation rate was observed in MPXV with 46 single nucleotide polymorphisms (SNPs) in the recent MPXV-2022 strains compared to the reference sequence from Nigeria-2022 (NC_063383.1).^13^ This necessitates complete genome sequencing of monkeypox cases to understand the evolutionary pattern of the circulating MPXV strain from the non endemic countries. Here, we report the findings of genomic characterization and phylogenetic analysis of ten confirmed monkeypox cases detected in India during July-August 2022 from Kerala (n=5 UAE travel history) and Delhi (n=5 without foreign travel history).

## Methods

During the period of July to August 2022, clinical specimens i.e., orophryngeal swab (OPS), nasopharyngeal swab (NPS), lesion crust and lesion fluids of 96 suspected Monkeypox cases were referred from 18 states (n=81) and 03 Union Territories (n=15) to ICMR-National Institute of Virology, Pune, India for diagnosis of Monkeypox. The clinical specimens of all the cases were tested using Monkeypox specific real time PCR.^14,15^ Of them, five cases each from Kerala and Delhi were found to be positive for MPXV. All the monkeypox negative cases (n=86) were also screened for Varicella zoster virus (VZV) and enterovirus (EV) specific real time PCR.^16,17^

The genomic characterization of these confirmed MPXV positive samples were carried out using next generation sequencing approach.^18^ Briefly, DNA was extracted from 200 μl of all the clinical specimens with commercially available DNA extraction kit (Qiagen, USA). The DNA concentration was measured by Qubit 2.0 fluorimeter using Qubit dsDNA kit (Themofisher Scientific, MA, USA). For preparation of libraries Nextera library kit from Illumina was used (Illumina, San Diego, CA, USA) following the manufacturer’s protocol. Using AMPure XP beads (Beckman Coulter, USA). The prepared libraries were quantified using the Qubit library quantification kit (Thermo Fisher, USA). The libraries were pooled and loaded onto Nextseq 2000 flow cell and sequenced on Illumina Nextseq2000 platform. The resulting FastQ sequencing files were imported and analyzed using CLC Genomics software 22.0.2 (Qiagen, USA).

The resulting raw data files were mapped with the NCBI MPXV reference sequence NC_063383 (MPXV-M5312_HM12_Rivers). The variant analysis was performed using CLC Genomics v22.0.2. Mutations having coverage more than 10X and frequency > 50 % was considered for the analysis. MPXV sequences of clade II, IIa, IIb, IIbA, IIb A.1, IIb A.2, IIb A.1.1 and IIb B.1 were further analyzed using MEGA v10. The nucleotide changes were further analyzed using Nextclade (https://master.clades.nextstrain.org).

For phylogenetic analysis seventy-five MPXV genome sequences (pre-outbreak sequences (n=21) as well as 2022 outbreak sequences (n=54) were downloaded from NCBI (n=72) and GISAID (n=3) database and were aligned with MPXV sequences from India. The alignment of sequences was done using MAFFT v7.505. The Maximum Likelihood (ML) tree was constructed using 1000 ultrafast bootstrap^19^ IQ TREE software^20^ version 2.2.0 with model finder.^21^

## Results

Of the 114 cases, MPXV infection was confirmed in ten cases from India using both *Orthopox* and Monkeypox specific real time PCR. Further the screening of monkeypox negative cases (n=86) indicated the presence of VZV (n=18) and EV (n=06) by real time PCR. The ten monkeypox confirmed cases were comprised of three male and two female from New Delhi (n=5) with no international travel history; while five males were from Kerala (n=5) with travel history from United Arab Emirates (UAE) to India^22^. All the cases were immunocompetent with no comorbidities (mean age: 31 years) and presented with short prodromal phase of fever, myalgia, vesiculo-pustular lesions primarily in genital area, face, trunk and extremities. Of all, nine cases had non tender firm lymphadenopathy in one or more sites (inguinal, cervical, submental, submandibular, retro-auricular), while one case didn’t show lymphadenopathy. All the cases have recovered without complications except a case from Kerala who succumbed to the infection following acute onset encephalitis^23^ (Table 1).

**Table 1:** Clinico-demographic, viral load and outcome of ten monkey pox cases from India, July-August 2022.

| Case No. | MCL No. | Age | Sex | Detected from | International travel history | Post Onset Dates | Sites of vesiculo-pustular lesion | Lymphadenopathy | Outcome | Monkeypox Viral Load (DNA copy/ml) | Percent Genome Recovered | Accession Nos. |
| --- | --- | --- | --- | --- | --- | --- | --- | --- | --- | --- | --- | --- |
| 1 | MCL-22-H-MPXV-29-5559 | 34 | M | New Delhi | No | 14 | Genitalia, bilateral palm, bilateral upper limb, scalp, trunk, and bilateral lower limb | No | Recovered without complications | 1.5×10<sup>9</sup> | 97.04 | EPI_ISL_15008575 |
| 2 | MCL-22-H-MPXV-52-5885 | 34 | M | New Delhi | No | 7 | Groin, leg, arm, trunk, face, scalp, palms, soles, and oral lesions | Inguinal Submental Retro-auricular | Recovered without complications | 1.15×10<sup>8</sup> | 99.91 | EPI_ISL_15022589 |
| 3 | MCL-22-H-MPXV-58-5982 | 35 | M | New Delhi | No | 10 | Groin, legs, arms, trunk, face, scalp, palm, sole | Inguinal and Submandibular | Recovered without complications | 9.13×10<sup>7</sup> | 98.67 | EPI_ISL_15022590 |
| 4 | MCL-22-H-MPXV-70-6050 | 31 | F | New Delhi | No | 5 | Bilateral arms, trunk, genitalia, face, scalp and mucosa of lower lip | Inguinal | Recovered without complications | 2.8×10<sup>7</sup> | 99.3 | EPI_ISL_15008576 |
| 5 | MCL-22-H-MPXV-84-6146 | 22 | F | New Delhi | No | 5 | Genitals, perianal, upper limb, trunk, face, lower limb and mucocutaneous lesions | Cervical and Inguinal | Recovered without complications | 1.3×10<sup>6</sup> | 99.94 | EPI_ISL_15008577 |
| 6 | MCL-22-H-MPXV-16-5316 | 35 | M | Kerala | UAE | 9 | Oral cavity, lips, genitalia, right infra-mammary, right tragus, right temple, and right deltoid region and hands | Cervical and Inguinal | Recovered without complications | 2.3×10<sup>8</sup> | 99.8 | EPS_ISL_13953610 |
| 7 | MCL-22-H-MPXV-21-5432 | 31 | M | Kerala | UAE | 9 | Both arms, genitalia, face, back, and neck | Cervical | Recovered without complications | 8.4×10<sup>8</sup> | 99.76 | EPS_ISL_13953611 |
| 8 | MCL-22-H-MPXV-28-5555 | 35 | M | Kerala | UAE | 8 | Face, neck, infra-axillary area, abdomen, fore-arm, left eye-lid, and back with carbuncle on right side of nape of neck | Cervical | Recovered without complications | 6.8×10<sup>8</sup> | 99.66 | EPI_ISL_14952916 |
| 9 | MCL-22-H-MPXV-51-5875 | 22 | M | Kerala | UAE | 16 | Single healed scrotal lesion with acute onset fever with encephalitis | Inguinal | Succumbed due to MPXV associated encephalitis | 1.75×10<sup>6</sup> | 91.2 | EPI_ISL_15008573 |
| 10 | MCL-22-H-MPXV-56-5969 | 30 | M | Kerala | UAE | 8 | Genitalia hands and face. | Inguinal | Recovered without complications | 2.2×10<sup>8</sup> | 94.11 | EPI_ISL_15008574 |

The MPXV genome (90 to 99%) could be retrieved from the skin lesions of confirmed monkeypox cases. All the retrieved MPXV sequences were aligned in MAFFT software version 7.505. The sequence alignment was checked manually in MEGA 10. The ML tree was constructed with a total of 85 sequences, 75 reference sequences downloaded from NCBI and GISAID database specific to MPXV Clade I, Clade IIa and Clade IIb along with ten sequences retrieved during current study.

The ML phylogenetic tree analysis placed the ten genome sequences from India (highlighted in blue) and eight genomes from USA (n=3), UK (n=2) and Thailand (n=3) under lineage A.2 clade IIb (Figure 1). Further they diverge into three sub clusters of A.2 lineage consist of total eighteen sequences. The MPXV sequences from India were grouped in two sub clusters; 7 sequences (Kerala n=5, Delhi n=2) aligned with the USA-2022 strain ON674051.1 and UK-2022 OP331335.1 formed first cluster. In this sub cluster, five sequences from Kerala were designated as A.2.1 based on the lineage defining mutations in the position C 25072 T, A 140492 C, C 179537 T. Two sequences from Delhi are lacking all three mutations hence still defined into A.2 lineage.

**Figure 1.**
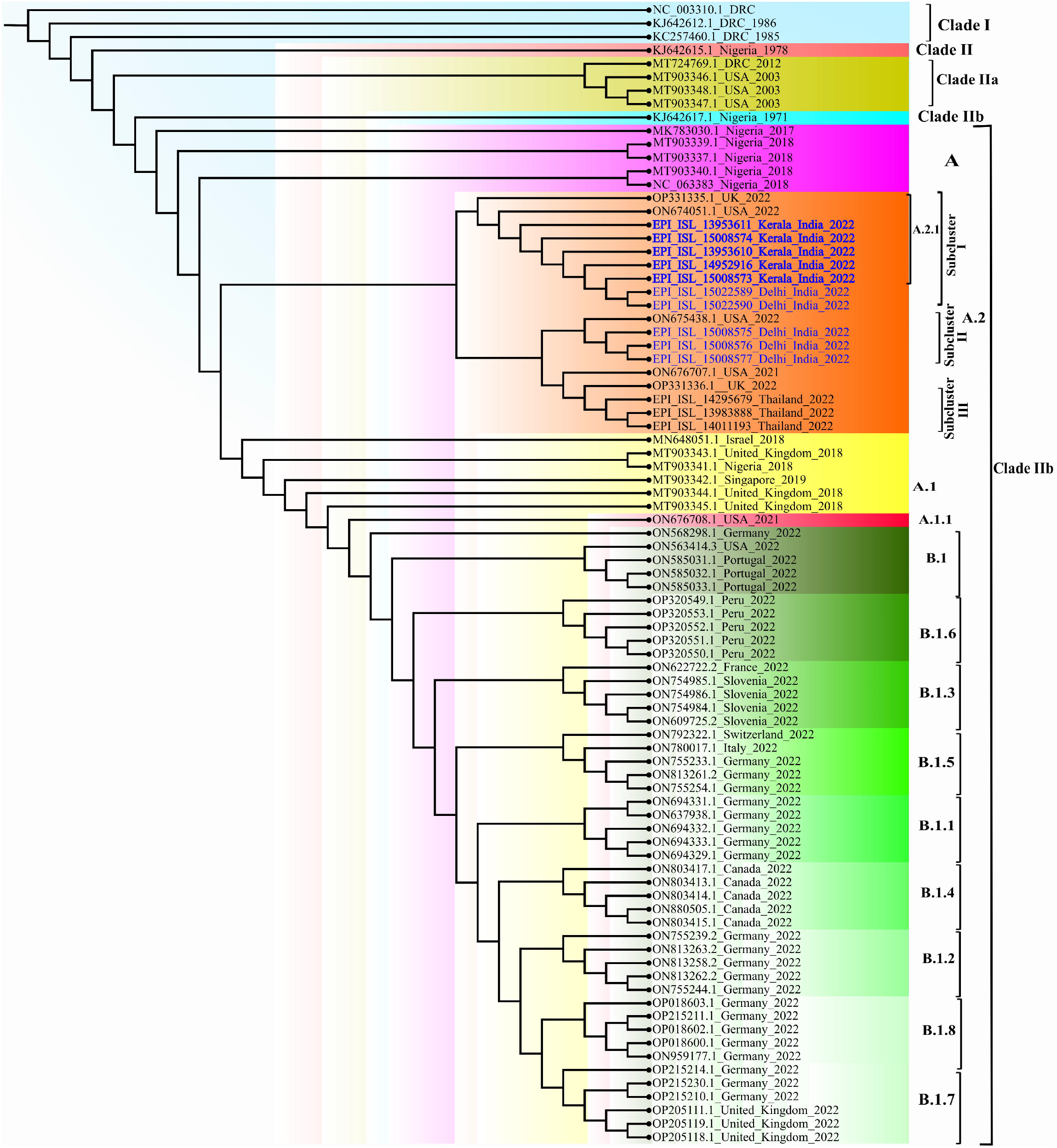
The maximum-likelihood phylogenetic tree of MPxV/hMPxV genome constructed using software IQ TREE with 1000 ultra bootstrap replication cycle. The retrieved sequences from ten monkeypox cases from India belong to A.2 lineage of Clade IIb (Highlighted in red color).

While it was noted that A.2.1 lineage defining mutations are lacking in the 3 sequences from second sub cluster of Delhi; aligned with USA-2022 strain ON675438.1. The third sub cluster consists of the MPXV sequences obtained from UK-2022 OP331336.1, USA-2021 ON676707.1 and three sequences of Thailand-2022 (Figure 1).

Analysis of Clusters of Orthologous Genes for orthopox viruses (OPG) revealed that the sub cluster I of seven sequences from India showed 07 synonymous mutations (OPG055, OPG071, OPG074, OPG093, OPG124, OPG163, OPG187) and 11 non-synonymous mutations (OPG040, OPG062, OPG063, OPG069, OPG074, OPG116, OPG160 and OPG208).

Sub cluster II included three sequences from India which indicated 04 non synonymous mutations (OPG025, OPG082, OPG084, OPG099) and 08 synonymous mutations (OPG037, OPG038, OPG105, OPG111, and OPG135). Mapping the A.2 sequences with the reference genome (NC_063383 strain) indicated total fifteen mutations (OPG031, OPG047, OPG053, OPG103, OPG113, OPG145, OPG174, OPG176, OPG180, OPG188, OPG190 and OPG205) which were common throughout all the three sub clusters of the A.2 lineage (Figure 2).

**Figure 2.**
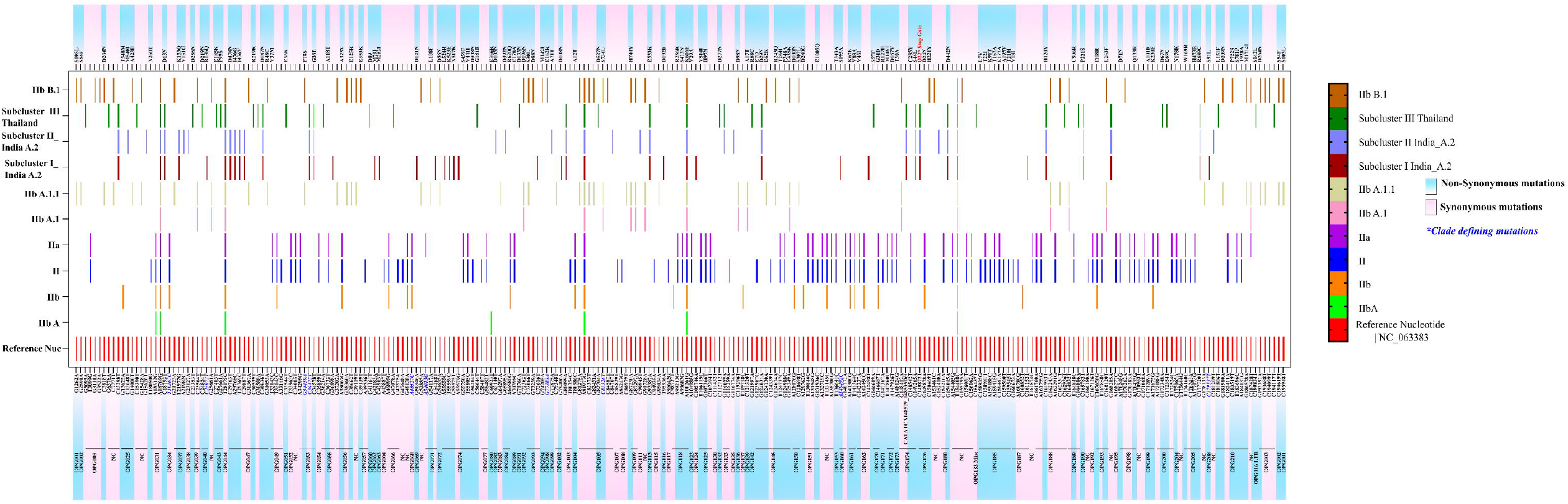
The variant analysis of MPXV genome was done using software MEGA 10 w.r.t reference sequence NC_063383. The depicted figure showed the synonymous and non-synonymous mutations in Clade II and lineages. The A.2 lineage is further characterized into sub-cluster I, II and III. All the clade defining mutations are marked with blue color positions.

We have also observed single nucleotide polymorphisms (SNPs) specific to sub cluster III compared to reference (NC 063383.1) which were not found in the sequences of sub cluster I and II. There were total 27 non synonymous changes observed in Thailand sequences.

APOBEC 3 mutation analysis indicated presence of 13 mutations in A.2 sequences from the current MPXV outbreak 2022. These could be A.2 lineage defining mutations apart from OPG053; C 34472 T reported during earlier studies (https://master.clades.nextstrain.org) (Figure 3). Further, we identified 25 additional APOBEC 3 mutations from the MPXV strain circulating in India. Besides this, 21 synonymous and non-synonymous APOBEC-3 mutations were also noted in sub cluster III.

**Figure 3.**
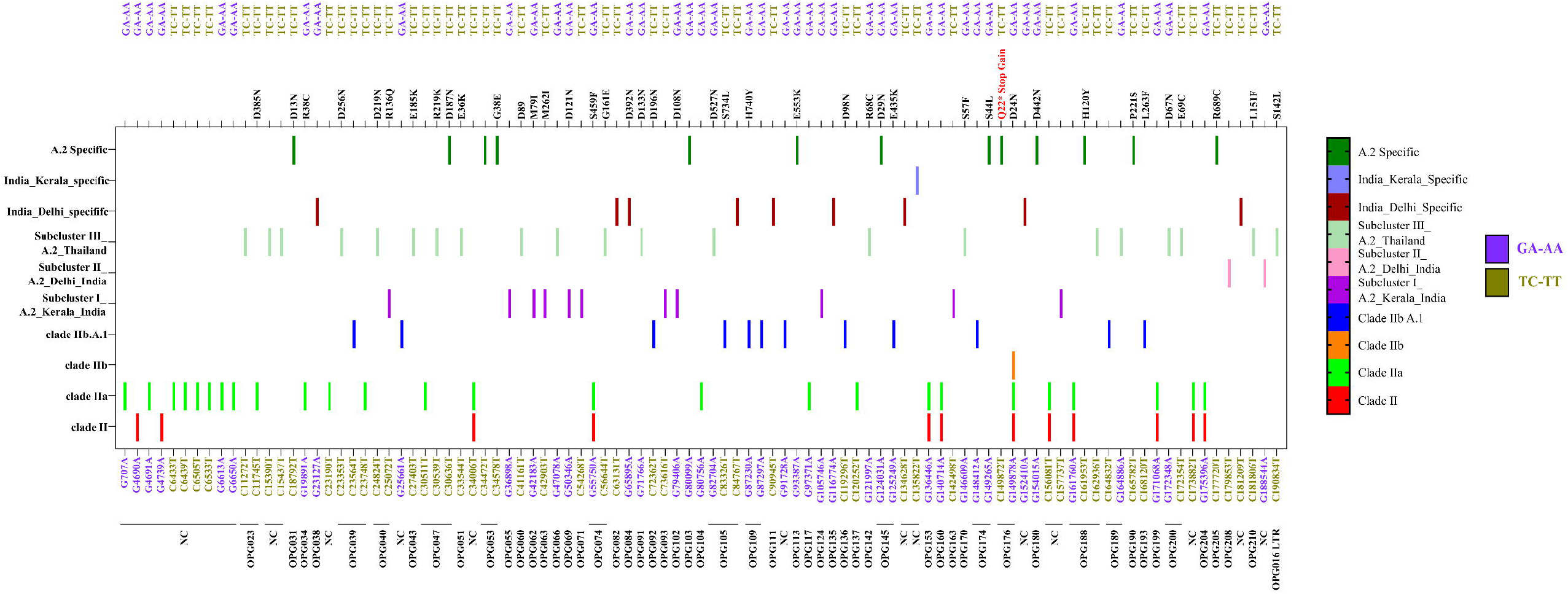
The APOBEC 3 mutation analysis was done using software MEGA 10.0. The mutations (GA-AA, TC-TT) were marked for all the clades.

Variant analysis of all sequences of A.2 lineage indicated a total of 34/67 synonymous mutations and 33/67 non synonymous mutations. Fourteen mutations are found in the non coding region and 53 mutations were observed in coding region of different ORFs. Most of the mutations are observed in gene OPG 047 which is closer to the middle of the genome followed by OPF 053, OPG 074 and OPG 105. Earlier reported clade defining mutation was observed (C 34472 T) gene on all the retrieved sequences from India. During complete genome analysis, we have observed insertions; one in OPG 047 (insertion of T at 29767), and other in non coding region (insertion of T at 170897). Also, a deletion was noted of CATATCA at (148529-148535) in gene OPG 174 ^20^ (Figure 4).

**Figure 4.**
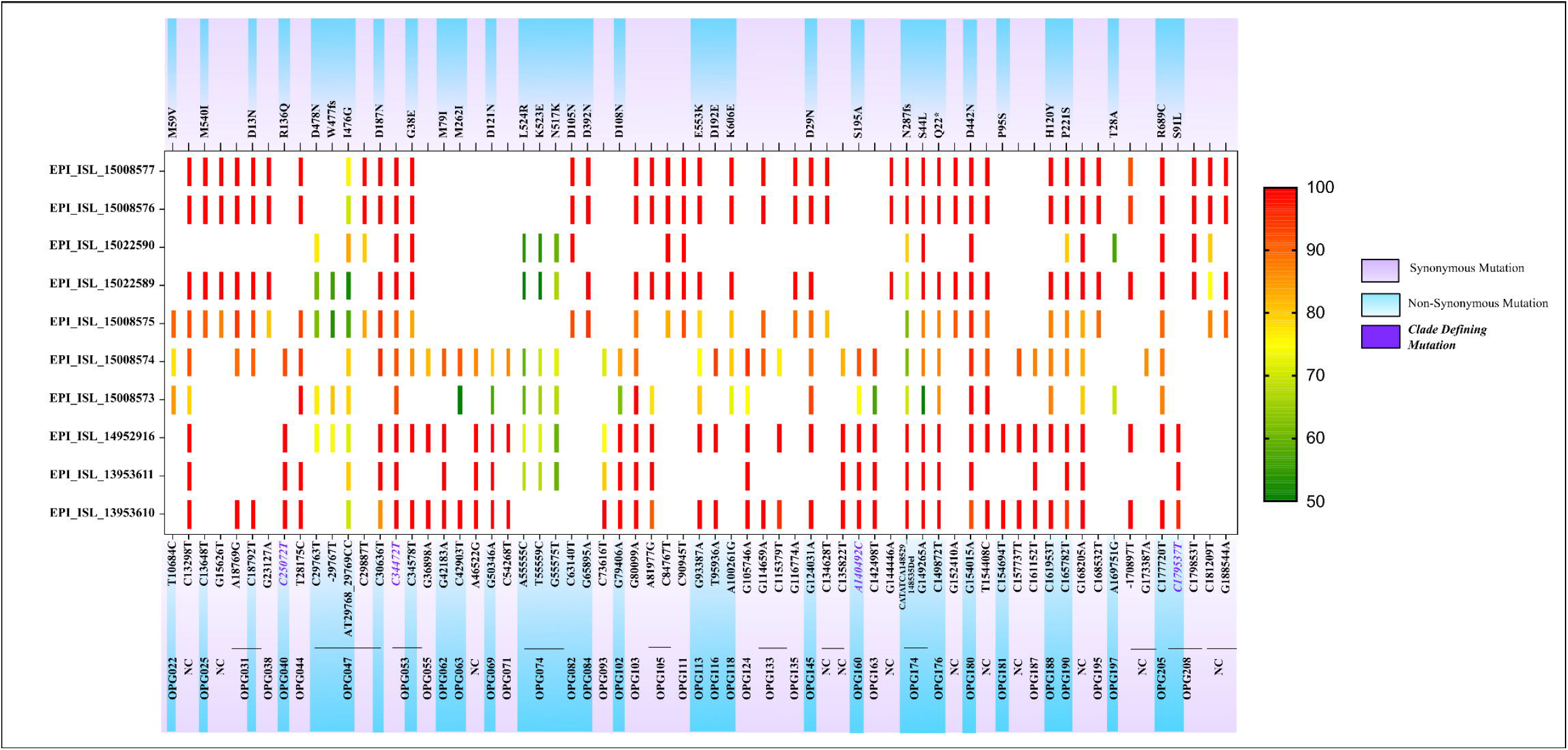
The variant analysis of MPXV genome of all the retrieved sequences was done using CLC genomics workbench with 50% frequency. The reference positions marked with synonymous and non-synonymous mutations for all the sequences with the respective frequencies.

Interestingly, one di-nucleotide substitution of AT-CC in OPG 047 at position 29768 leading to amino acid change (I 476 G) was observed in the retrieved sequences of ten monkeypox cases. Of 67, a total 63 substitution mutations were observed in cluster from India in which 58 are transitions and 05 are transversion (Non synonymous 30 and synonymous 33). One substitution mutation which led to stop gain at position 149872 was also recorded in gene OPG 176. No mutations were noted on H3L (OPG 108), glycosil transferase which plays important role in pox virus entry into host cell. We have also analyzed MPXV gene OPG 31 and OPG 174 that are predicted to modulate the host immune response and observed four non synonymous and two synonymous SNPs.

## Discussion

The outbreak of MPXV among non-endemic countries raised concerns and emphasized to prepare for diagnosis and genomic sequencing to understand its linkage. India started preparing its virus research diagnostic laboratory network for augmenting diagnosis capacity for MPXV. The preparedness to deal with emerging situation of monkeypox, a high alertness and intensive risk communication were directed at identified sites in healthcare facilities (skin, pediatric, immunization clinic and intervention sites identified by National AIDS Control Organization for men having sex with men and female sex workers). India quickly identified the first two cases of monkeypox among UAE returnees during July 2022. The genomic analysis revealed the infection of these cases with lineage A.2 of sub clade IIb.^22^ Gigante et al., demonstrated 80 nucleotide changes in lineage A.2 compared to the B.1 lineage which has been predominant lineage of 2022 suggesting an independent virus strain emergence.^23^ The genomic research on the 2022 MPXV outbreak has also grabbed the attention of the researchers with divergence of lineage B.1 from A.1 lineage of 2018-2019 outbreaks.^24^

The 2022 Monkeypox outbreak demonstrates the accelerated microevolution of MPXV leading to the divergence in the viral phylogeny. As on 6 September 2022, a total of 864 MPXV genome sequences were available in NCBI and GISAID databases consisting of Clade I (n=30), Clade II (n=01), Clade IIa (n=17), Clade IIb A (n=12), Clade IIb A.1 (n=06), Clade IIb A.2 (n=08), Clade IIb A.1.1 (n=01), Clade IIb B.1 (n=476), IIb B.1.1 (n=102), IIb B.1.2 (n=61), IIb B.1.3 (n=35), IIb B.1.4 (n=22), IIb B.1.5 (n=16), IIb B.1.6 (n=10), IIb B.1.7 (n=45), IIb B.1.8 (n=22). Despite this, there is still scarcity of genomic data on lineage A.2 with availability of only 8 genome sequences excluding the sequences from India.

Here, we presented the genomic and phylogenetic analysis data of ten monkeypox cases detected in India. All the MPXV genome sequences retrieved from these cases belonged to lineage A.2 which further diverged into three sub clusters. The seven sequences grouped into sub cluster I showing highest similarity with MPXV_USA_2022_FL001 also called as A.2.1, while sub cluster II having three sequences aligned with USA_2022_VA001 sequences of lineage A.2 clade II b reported from USA 2022. Delhi MPXV sequences in sub cluster I and II are showing divergence which needs to be further explored.

The sub cluster III has monkeypox sequences reported from UK, Thailand during current outbreak of 2022 and in USA during 2021. These sequences have many shared mutations that separated them from other two subcluster including India (A.2) and other travel-associated cases from 2017–2021 of other lineages (A, A.1, A.1.1). The A.2 lineage MPXV sequences from India showed a divergence from the MPXV sequences reported from Germany, Italy, Portugal, Switzerland and France (lineage B.1) and earlier outbreak sequences from Nigeria, Israel and Singapore 2017/18 (lineage A.1). The findings of our study are in concordance with the recent study of Gigante et al, which reported the circulation of both A.2 and B.1 lineages in current outbreak in USA with similarity to MPXV sequences of traveler from Nigeria to Texas in 2021.^23^

Recently O’Toole and Rambaut reported that the APOBEC3 with cytidine deaminase activity could be significant factor in the short-term evolution of MPXV since 2017.^27^

Although it is helpful to know what is driving the mutational changes, it is equally crucial to understand how genetic alterations are facilitating human-to-human transmission. Our analysis of mutations in APOBEC3 in the current circulating Clade IIb supported a strong inclination for GA to AA and TC to TT mutations, indicating cytosine deaminase functioning reported only in Clade II since 2017 and not in Clade I.^25^ Our data revealed additional 13 APOBEC3 mutation and 16 SNPs in all the available sequences from the current MPXV outbreak 2022 which might contribute as lineage defining changes in the A.2 lineage. A recent study Isidro et al. also demonstrated microevolution and divergence of MPXV sequences of 2022 outbreak pertaining to APOBEC3 and other proteins.^24^

Genomic evolutions of the Orthopoxviruses such as MPXV might increase the possibility of higher transmission and host range which can affect larger population. Overall poxviruses have shown slower but the high recombination in genome leading to gene gain or loss due to mainly selective pressures. The *Orthopoxvirus* genes (OPGs) are composed of about 100 genes that encode proteins involved in the process of virus reproduction i.e., virion structure, replication, and transcription and about 50 genes from both ends that encode accessory proteins responsible for virus-host interactions.^26^ An analysis of OPG genes indicated nucleotide changes in 24 OPGs between sub cluster I and II leading to insertions, deletions or substitutions in various important gene, while the analysis of sub cluster III indicated mutations in 22 OPGs. We have also noted a seven-nucleotide deletion in all three A.2 sub clusters in OPG 174 gene. It is known to play role in influencing virulence by suppressing immune system; further studies are needed to determine the impact of this deletion on virulence of A.2 lineage.

A highly conserved 120Kb of core region of *Orthopoxvirus* is thought to code for basic viral functions and the terminal region genes responsible for host adaption.^27^ The changes during host adaption may play important role in disease severity as well as spread.^27^ Hence, genome evolution mechanisms and importance of gene functions needs to be studied further to understand evolution of the MPXV genome. As the B.1 is found to be the predominant lineage of 2022 Monkeypox outbreak globally, the introduction event of A.2 lineage specifically in the USA, UK, India and Thailand is the question of further exploration.

## Ethical approval

The study was approved by the Institutional Human Ethics Committee of ICMR-NIV, Pune, India under the project ‘Providing diagnostic support for referred samples of viral hemorrhagic fever and other unknown etiology and outbreak investigation’. The clinical data collected were anonymized. The informed consents were obtained for the use of the clinical details in the study.

## Author Contributions

PDY, AMS contributed to study design, data analysis, interpretation and writing and critical review. AK, SP, DYP, YJ, TM, VR, RRS, MV, PG, AV, ArK, SD, ABK, SC, SK contributed to sample collection, data collection, interpretation, writing and critical review. PDY, AMS, DYP, RRS, AK, PA contributed to the critical review and finalization of the paper.

## Acknowledgement

Authors extend gratitude to Smt. Veena George [Hon’ble Minister for Health and Family Welfare, Kerala], for efficient coordination of the monkeypox virus disease control activities, and “The Team Kerala Health,” the district administration. We would also extend our gratitude towards Dr. Asha Thomas, Additional Chief secretary of Medical Education, Mrs. Tinku Biswal, Principal Secretary for Health Kerala, Dr. Thomas Mathew, Director of Medical Education and Dr. Preetha PP, Director of Health Services, Kerala. The authors would like to thank Dr. Lakshmi Geetha Gopalkrishnan, State Epidemiologist, and District Surveillance Officers of Thiruvananthapuram [Dr. Preethi James], Kannur [Dr. Shaj MK], Kollam [Dr. Sandhya], and Thrissur [Dr. Anoop TK and Dr. Kavya Karunakaran]. Dr. Kala Kesavan P, Principal; Dr. Nizarudeen A, Medical superintendent; Dr. Aravind Reghukumar, HOD Infectious Diseases and Dr. Manjusree Suresh, from Government Medical College Thiruvananthapuram. The authors are thankful to Dr. Prathap Somanath, Principal, Dr. Sudeep, HOD Infectious Diseases; Dr. Manasi Ravindranath from Government Medical College Kannur; Dr. Shinas, Government Medical College, Manjeri. The authors also acknowledge the support from Dr. Fazil Abubaker from Daya General Hospital and Specialty Surgical Centre, Thrissur.

Authors extend sincere thanks to Dr. Bijayalaxmi Sahoo, Professor and Head, and resident doctors Dr. Abhinav Kumar, Dr. Aneet Kaur, Dr. Bhawna Solanki, Dr. Anjali Bagrodia of Dermatology; Dr. Sonal Saxena, Professor, and Head, Department of Microbiology from Maulana Azad Medical College and Lok Nayak Hospital, New Delhi for providing support for sample collection and transportation. We are also grateful to Dr. Lalit Dar, Professor, Department of Microbiology; Dr. Aashish Choudhary, Dr. Megha Brijwal, from All India Institute of Medical Sciences, New Delhi. The authors are thankful to Dr. Avdesh Kumar, State Surveillance Officer, New Delhi and his team for coordination. The authors are extremely grateful to Dr. Nivedita Gupta, Scientist ‘F’ and Head, Epidemiology and Communicable Diseases, ICMR, New Delhi for her constant support.

We also acknowledge the excellent technical support from Dr. Kannan Sabarinath PS, Dr. Rajlaxmi Jain, Ms. Jyoti Yemul, Mr. Sunil Shelkande, Ms. Pratiksha Vedhpathak, Mrs. Shubhangi Sathe, Ms. Vaishnavi Kumari, Ms. Nandini Shende, Mr. Raj Hawale for the diagnosis and data management for the diagnosis and data management. The authors also would like to thank and express immense gratitude to the monkeypox cases and family members, who willingly agreed and provided consent to be part of the study.

